# Census: accurate, automated, deep, fast, and hierarchical scRNA-seq cell-type annotation

**DOI:** 10.1101/2022.10.19.512926

**Authors:** Bassel Ghaddar, Subhajyoti De

**Affiliations:** Center for Systems and Computational Biology, Rutgers Cancer Institute of New Jersey, Rutgers University; 195 Albany St., New Brunswick, New Jersey 08901

## Abstract

We developed Census, an automated, hierarchical cell-type identification method for scRNA-seq data that can deeply annotate normal cells in mammalian tissues and identify malignant cells and their likely cell of origin. When benchmarked on 44 atlas-scale normal and cancer, human and mouse tissues, Census significantly outperforms state-of-the-art methods across multiple metrics. Census is a fast and fully automated method, although users can seamlessly train their own models for customized applications.

## Main text

Single cell RNA-seq (scRNA-seq) has enabled annotation and transcriptional characterization of cell-types in multicellular species. Cell-type annotation is a critical and often difficult and time-consuming first step in scRNA-seq data analysis. Typical annotation pipelines involve cell clustering followed by comparison of cluster differentially expressed genes with celltype marker genes databases^1^. While this approach is suitable for major, well-defined cell-types, it can be challenging to annotate cells from noisy datasets or to identify cell-subtypes for which marker genes are overlapping, poorly expressed, or incompletely described^2,3^. This problem is especially pertinent while analyzing scRNA-seq data from perturbation experiments, disease contexts such as cancer, or treatment conditions.

As such, a number of automated cell identification methods have been developed^4–13^, and cell types in many organ-types have been annotated^14,15^. However, when applied to complex tissues, in practice, most suffer from a number of limitations^16–18^. These include inaccurate or shallow annotations, limited organ or cell-type scope, long computation time, the requirement of large reference data, or an inability to distinguish between malignant cells and their normal counterparts^16–18^. In addition, batch effects or differences in cell subtypes between reference and test data often lead to incorrect label predictions or resolutions. Without clearly defined hierarchical cell-type relationships, it can be difficult to identify the appropriate label resolution or alternative cell-type annotations.

To overcome these limitations we developed Census, a fast and fully automated hierarchical cell-type identification method that is conceptually motivated by inherently stratified developmental programs of cellular differentiation. Census implements a collection of hierarchically organized gradient-boosted decision tree models^19^ that successively classify individual cells according to a predefined cell hierarchy (**Fig. 1a**). Briefly, Census begins by identifying a cell-type hierarchy from reference scRNA-seq data by hierarchically clustering pseudo-bulk cell-type gene expression data using Ward’s method, which splits each node into two child nodes. Next, starting with the root node and for each successive node, differentially expressed genes that distinguish cells from the two child nodes are identified and used as features to train a gradient-boosted tree model to classify the node identity of individual cells. Census uses multiple, relevant percentile-ranked feature scores, allows for missing values, and trains on both full and sparsely down-sampled data, resulting in models that are robust to batch effects.

**Figure 1.**
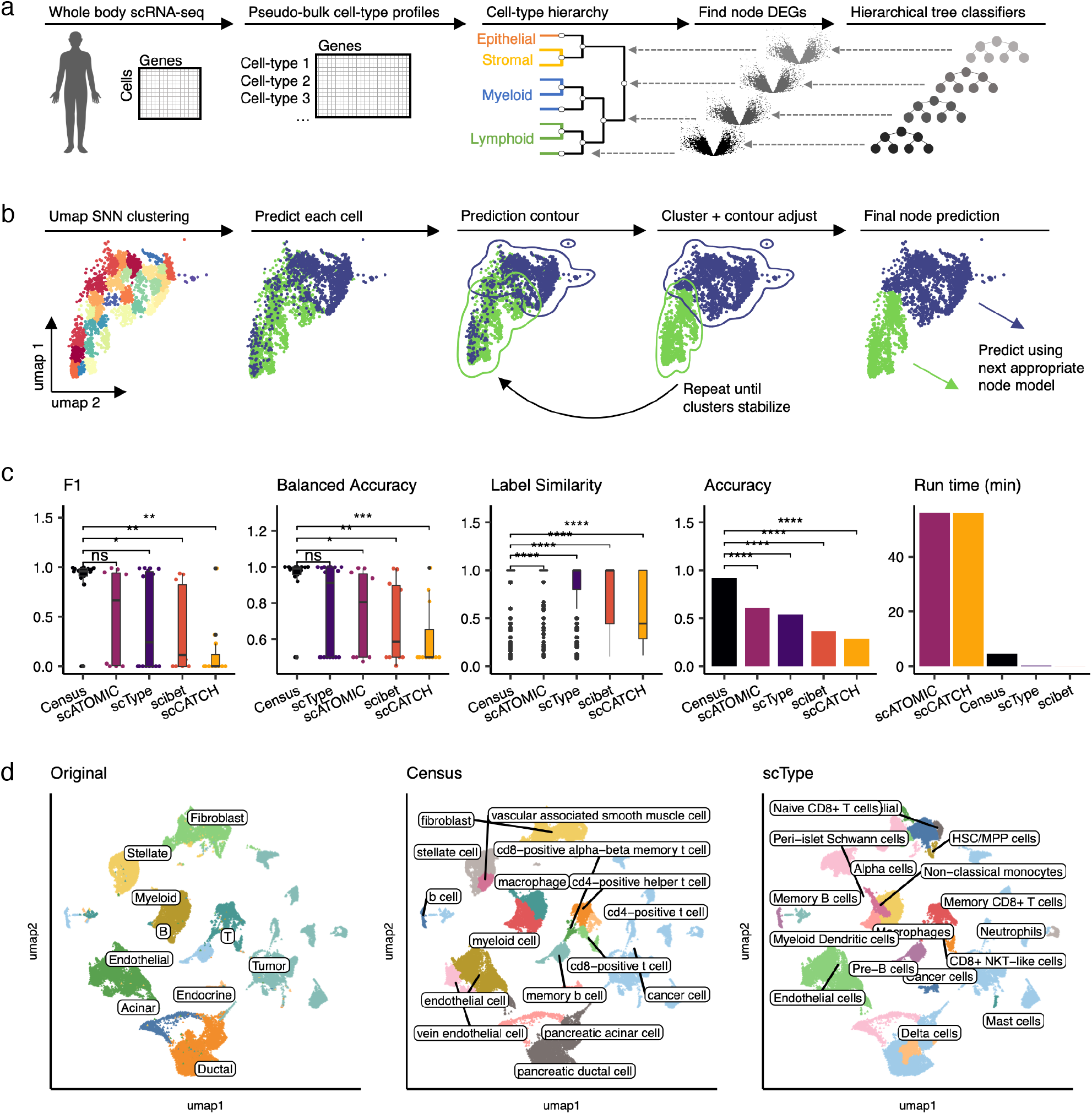
Overview of Census and initial benchmarking. **a,** Schematic diagram of training a Census model. The default Census model is trained on the Tabula Sapiens and Cancer Cell Line Encyclopedia. **b,** Schematic diagram of the Census prediction method and label-stabilizing algorithm. **c,** Benchmarking Census against four other automated cell-type annotation methods using a pancreatic cancer dataset. Boxplots show median (line), 25^th^ and 75^th^ percentiles (box) and 1.5xlQR (whiskers). Points represent outliers; Wilcoxon tests, **** p<1e-4; *** p<1e-3; ** p<1e-2, * p<0.05; ns, not significant. **d,** Uniform manifold approximation and projection (UMAP) plots of 57,530 cells from a pancreatic cancer study colored by cell-type with overlaid labels. Left, original study annotations, middle, Census annotations, right, scType annotations. HSC/MPP, hematopoietic/multipotent progenitor.

New datasets are annotated using the pretrained models followed by a custom developed label-stabilizing algorithm (**Fig. 1b**). Census first uses uniform manifold projection and approximation (UMAP) and a shared nearest-neighbor graph^20^ to finely cluster data, and it begins by annotating cells with the root classifier. Next, the average label per cluster is propagated and prediction contours in UMAP space are computed. Census resolves disputes within overlapping contour regions and repeatedly redraws contours until the prediction contours stabilize. Given each cell’s new identity, the next appropriate node classifier is applied, and this process is repeated until terminal classifications are reached. This is particularly advantageous for organ-scale data with many cell subtypes, both in terms of computation time and classification performance. Census thus leverages multiple design features to achieve high speed and accuracy.

We trained Census to classify 175 cell-types from 24 organs using data from the Tabula Sapiens^14^. Construction of the cell-type hierarchy revealed biologically meaningful groups, with the largest split being immune vs. non-immune cells and with cells further segregating into lymphoid, myeloid, endothelial, stromal, and epithelial groups (**Fig. S1a**). To identify cancer cells, we trained models on scRNA-seq data from 19 cancer types from the Cancer Cell Line Encyclopedia^21^ to distinguish malignant cells from organ-specific normal epithelium. The total collection of models had 351 nodes, and all node models had high training classification accuracy (median AUC=0.99, **Fig. S1b**).

We first benchmarked Census against four other state-of-the-art automated annotation methods (scType^4^, scATOMIC^5^, scibet^6^, scCATCH^7^) using a pancreatic cancer dataset^22^ that included 57,530 normal and malignant epithelial, stromal, and immune cells. Annotation performance was evaluated by five metrics: F1 score, balanced accuracy, total accuracy, run time, and “label similarity” scores that we computed using our predefined cell hierarchy to quantify closeness of the predicted label to the study’s original annotation (see Methods). Census was the top performing method with regards to prediction quality, where it had a higher mean F1 score and balanced accuracy than the second-place method and significantly higher scores than the others (Wilcoxon, p<0.05), and it had significantly higher label similarity and accuracy than all methods (Wilcoxon, p<2e-16, **Fig. 1c**). While Census was not the fastest method, it ran in 4.5 minutes (range: 1 second-56 minutes, **Fig. 1c**). Census correctly identified 9/10 major cell-types, distinguished between normal and malignant epithelial cells, and identified deeper immune subtypes than originally annotated (**Fig. 1d**).

In terms of accuracy, speed, and precision, Census and scType were the top two methods (**Fig. 1d, Fig. S1c**). We thus proceeded to assess their annotation performance on 44 other challenging normal and cancer datasets from human tissues and from the Tabula Muris^15^; these data included 1,769,071 total cells from 105 harmonized cell labels. In aggregate, Census had significantly higher F1 scores, balanced accuracies, label similarities, and overall accuracies than scType (all Wilcoxon p<2e-16, **Fig. 2a**), although scType had shorter run times (Wilcoxon p<2e-16, **Fig. 2a**) - though Census was still very fast with an average annotation speed of 13,000 cells/minute. Looking at prediction performance in individual studies, Census had higher mean values than scType in 83/100 commonly evaluable metrics (**Fig. 2b**). These data place Census as a top automated annotation method (**Table S1**).

**Figure 2.**
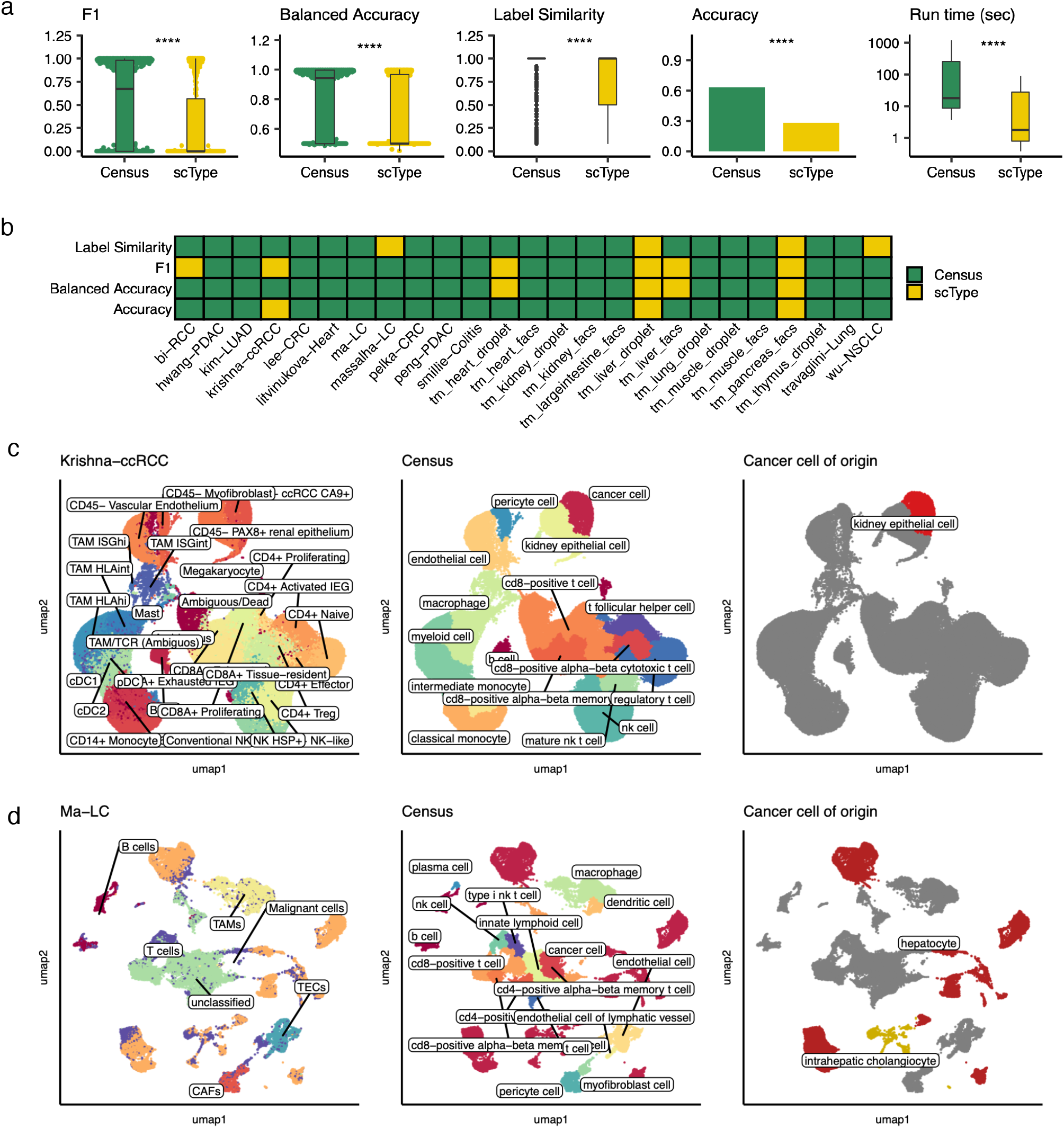
Extended benchmarking of Census on normal and cancer, human and mouse tissues. **a,** Performance metrics comparing Census and scType on 25 tissues. Boxplots show median (line), 25^th^ and 75^th^ percentiles (box) and 1.5xlQR (whiskers). Points represent outliers; Wilcoxon tests, **** p<1e-4. **b,** Heatmap showing the top performing method across four evaluation metrics for 25 commonly evaluable tissues. Green color, Census had a higher mean value, yellow color, scType had a higher mean value. tm, Tabula Muris **c,** Example uniform manifold approximation and projection (UMAP) plots of 167,283 cells from a clear cell renal cell carcinoma (ccRCC) study colored by cell-type with overlaid labels. Left, original study annotations, middle, Census annotations, right, Census cancer cell of origin prediction. TAM, tumor associated macrophage; DC, dendritic cell, Treg; regulatory T-cell. **d,** Example UMAP plots of 57,000 cells from a liver cancer (LC) study colored by cell-type with overlaid labels. Left, original study annotations, middle, Census annotations, right, Census cancer cell of origin prediction. TAM, tumor associated macrophage; TEC, tumor endothelial cell, CAF, cancer associated fibroblast.

Inspection of specific results highlights the power of Census in diverse settings. In two datasets each of human breast^23,24^, colon^25,26^, kidney^27,28^, liver^29,30^, lung^31,32^, and pancreas^22,33^ cancers, Census identified malignant cells and distinguished them from concurrent normal epithelium (**Fig. 2c–d, Fig. S2a**). It also identified the likely cell of origin for cancer cells. For example, in liver cancer, Census correctly distinguished between known hepatocellular vs. cholangiocarcinoma cells (**Fig. 2d**), in pancreatic cancer it identified most malignant cells as ductal cells and a few tumors as having an acinar cell origin, and in colon cancers the cells of origin were from the enterocyte lineage (**Fig. S2a**). On deeply annotated normal tissue atlases, Census distinguished between several cell subtypes. For example, it identified aerocytes and capillary, vein, artery, and lymphatic endothelial cells, distinguished between alveolar, adventitial, and myo-fibroblasts, and identified several T-cell and myeloid cell subsets in lung^34^, colon^35^, and heart^36^ tissues (**Fig. S2b**). Census also had excellent performance on mouse tissues when tested on droplet and plate-based sequencing samples from the Tabula Muris^15^ (**Fig. S2c**), with a mean balanced accuracy of 0.8, label similarity of 0.89, and run time of 13 seconds across all tissues. Overall, Census correctly identified 81/105 tested cell subtypes (compared to 35/89 by scType), and Census’s prediction accuracy per cell-type correlated with the corresponding number of cells used for model training (Spearman p=0.26, p=0.01).

In summary, Census enables easy and fast, fully-automated cell-type identification from scRNA-seq data using a hierarchical cell-type reference. It significantly outperforms other state-of-the-art methods when extensively tested on human and mouse, cancer and normal tissues. Utilization of a cell-type hierarchy provides a natural interpretation of annotation results and aids in Census’s superior performance. It also allows cell type identification at different resolutions, which can be advantageous when comparing results from different datasets, across species, and in the context of complex diseases such as cancer. While the core Census model is trained on the Tabula Sapiens^14^, with one line of code users can seamlessly train their own models with other references for customized applications. Census is available on our Github: https://github.com/sjdlabgroup/Census.

## Methods

### Census algorithm

#### Constructing the cell-type hierarchy

Census begins by constructing a cell-type hierarchy from reference scRNA-seq data. Given all gene expression and cell-type labels, pseudo-bulk celltype profiles are created by summing gene counts across all barcodes per cell-type, creating a gene by cell-type table. The resulting profiles are TP10K normalized and then hierarchically clustered using Ward’s method, which clusters each node into two leaves. Each node of the hierarchical tree is numbered, and the terminal leaves represent the final cell-types.

#### Training a Census model

A collection of gradient-boosted tree-based classification models^19^ organized by the cell-type hierarchy are next trained to predict cell-type label from scRNA-seq gene expression values. Each node of the cell-type hierarchy has an associated classification model; there are as many models as there are nodes in the cell-type hierarchy. Starting with the root node of the cell-type hierarchy, the cells of all downstream cell-types whose lineage contains the given node are gathered. All nodes bifurcate into two child nodes; the task of the node model of the given node is to classify cells into the appropriate child node of the given node. Cells from the training data are thus given the new identity of their respective child node of the given node through which their lineage runs. This results in two identity classes, and marker genes that distinguish these two classes are identified using Wilcoxon Rank Sum testing, as implemented in Seurat^20^. By default, all statistically significant marker genes are used, although users may impose custom filters or provide alternate marker gene data. The node model uses marker gene counts data to predict the associated cell-label.

Census modifies the training data in three ways before model training. First, zero-values are replaced with NA (not available) to be treated as missing values by the classification algorithm. This is done to account for variable dropout levels across scRNA-seq datasets and across individual cells in a given dataset, wherein zero-values may represent lack of detectable gene expression, low gene expression confounded by measurement noise, or uncaptured gene expression. This also accounts for potentially missing genes between the training data and test datasets, and it takes advantage of the underlying sparsity-aware split finding algorithm developed explicitly by xgboost^19^ to optimize handling of missing values in classification problems. Second, gene values for each cell, excluding missing values, are percentile ranked. Third, Census sparsely down-samples this data to create additional training data with more missing values; the default is to supplement the full training data with a 90% sparsified dataset, i.e. a dataset with 90% of values replaced with missing values. The full and sparse training data are combined, and a classification model is trained to predict the cell-label given the gene expression data. This process is repeated for each node of the cell-type hierarchy, with the final models predicting terminal cell-type labels. The above design choices make Census robust to missing values and batch effects.

#### Annotating cell-types in new datasets

Census uses the resulting models in conjunction with a custom label-stabilizing algorithm to predict new datasets. First, the test dataset is processed using standard scRNA-seq pipelines to project it in two dimensions using uniform manifold projection and approximation (UMAP, i.e. by TP10K normalization, scaling, finding variable genes, computing principle components, and then running the UMAP algorithm using the top principle components), as implemented in Seurat^20^. This data is finely clustered using the first two UMAP dimensions using a shared nearest-neighbor (SNN) algorithm, as implemented in Seurat^20^. These clusters represent groups of highly similar cells in the test dataset and are used to mitigate prediction error in individual cells. It is crucial at this step that high resolution clustering is done to take advantage of UMAP’s preservation of local structure and to avoid co-clustering distant cells.

Next, starting with the first model corresponding to the root node of the cell-type hierarchy, new cell identities are predicted for each individual cell in the test dataset. Census then implements a custom label-stabilizing algorithm that counteracts potential dataset noise and prediction error. First, the average label is propagated within each UMAP SNN cluster. Next, prediction contours are computed on the UMAP plot using the MASS^37^ R package. In areas where prediction contours do not overlap, all cells within the contour are given the identity of the contour. In areas where the prediction contours overlap, cells within the overlapping region are given the identity of the most common label in that region. After resolving contour disputes, the most common label is again propagated across each UMAP SNN cluster, and new prediction contours are computed. This process is repeated until either there are no more overlapping prediction contours or until there are no further changes to any cell labels. Each cell now has a new identity, and the next appropriate node model is used to predict subsequent labels; this process is repeated until terminal cell-type classifications are reached. A record of predicted classes and probabilities for each cell in each round of classification is retained.

### Census models

The core census model was trained on the Tabula Sapiens^14^ to classify 175 cell-types from 24 organs. The cell-type hierarchy contained 345 nodes, with the first node bifurcating into immune vs non-immune cells, and then further branches dividing into B-lymphoid, T-lymphoid, myeloid, endothelial, stromal, and epithelial compartments. The Tabula Sapiens was chosen as the reference for the core model due to its comprehensive human body profiling and consistent cell class ontology labeling, and through extensive benchmarking experiments the core model was found to generalize well across a range of datasets. To predict cancer cells, we trained models on scRNA-seq data from 19 cancer types from the Cancer Cell Line Encyclopedia^21^ to distinguish malignant cells from organ-specific normal epithelium. For example, to identify cancer cells in the pancreas, the classification model was trained to distinguish between the cancer cell line data and pancreas epithelium from the Tabula Sapiens, i.e. ductal, acinar, and endocrine cells. When predicting new datasets, the Census model begins by finding terminal classifications for all celltypes using the Tabula Sapiens trained model and cell-type hierarchy. If cancer cells are expected in the sample, then the organ-specific cancer model is applied only to the cells classified as epithelial cells by the Tabula Sapiens model to identify cancer cells. The same contour and cluster-based label stabilizing algorithm is applied. The final output will contain cell-type predictions, and for the predicted cancer cells, and it will also retain the origin normal cell-type prediction as the predicted cell of origin. Cancer cell type models are available for the following organs: breast, colon, kidney, liver, lung, and pancreas. While the Tabula Sapiens and cancer models enable rapid and automated cell-type identification for a variety of datasets, users can also easily train new models with other references (which may include cancer cells as part of the reference) for custom applications.

### Benchmarking analyses

Initial benchmarking compared Census to four other state-of-the-art automated annotation methods: scType^4^, scATOMIC^5^, scibet^6^, and scCATCH^7^. For scType, the primary tissue type as well as ‘immune system’ were chosen for as the tissue type. For scibet, the “30_major_human_cell_types” model was used. Other methods were run with default parameters. All methods were evaluated using on a pancreatic cancer dataset^22^. Census and scType were further evaluated on two datasets each of colon^25,26^, kidney^27,28^, liver^29,30^, lung^31,32^, and pancreas^22,33^ cancers, normal lung^34^, colon^35^, and heart^36^ datasets, and tissues from the Tabula Muris^15^ where applicable. Census was additionally evaluated on two datasets of human breast cancer^23,24^. In total, Census was evaluated on 44 tissue samples from 23 unique tissue types that contained 1,769,071 total cells from 105 harmonized cell labels. To assess performance, F1 scores and balanced accuracies were calculated using the caret R package (https://topepo.github.io/caret/index.html). Label similarity scores to assess closeness of a predicted label to the original author annotated label were calculated as follows. First, cell-type labels from the original studies and the predicted labels from scType^4^, scATOMIC^5^, scibet^6^, and scCATCH^7^ were harmonized to the cell ontology annotations used in the Tabula Sapiens^14^ using the closest matching label. Then using the cell-type hierarchy created from the Tabula Sapiens, the label similarity score was calculated for each cell-type prediction as the percent of shared nodes of the shorter of the lineages of either the author annotated label or the predicted cell-label. Each individual cell thus had a label-similarity score, and each cell-type from each tissue sample had an F1 score and balanced accuracy. Wilcoxon Rank Sum tests were used to compare metrics for the different cell-type annotation methods.

## Statistical analyses

All statistical analyses were performed using R version 3.6.1 (https://www.r-project.org/). The ggpubr package (https://github.com/kassambara/ggpubr) was used to compare group means with nonparametric tests. P-values reported as <2e-16 result from reaching the calculation limit for native R statistical test functions and indicate values below this number, not a range of values. Data processing relied heavily on the Tidyverse v1.3.2 R packages (https://www.tidyverse.org/).

## Supplementary Materials

**Table S1.** Evaluation metrics for benchmarking experiments

## Acknowledgments

We acknowledge the Office of Advanced Research Computing (OARC) at Rutgers, The State University of New Jersey for providing access to the Amarel cluster URL: https://it.rutgers.edu/oarc.

## Author contributions

BG conceived the study. BG designed and performed all data analyses. BG and SD interpreted the data and wrote and revised the manuscript.

## Data and materials availability

Census is available on our Github: https://github.com/sjdlabgroup/Census. Other data and code are available from the authors upon reasonable request.

---

**Figure S1.**
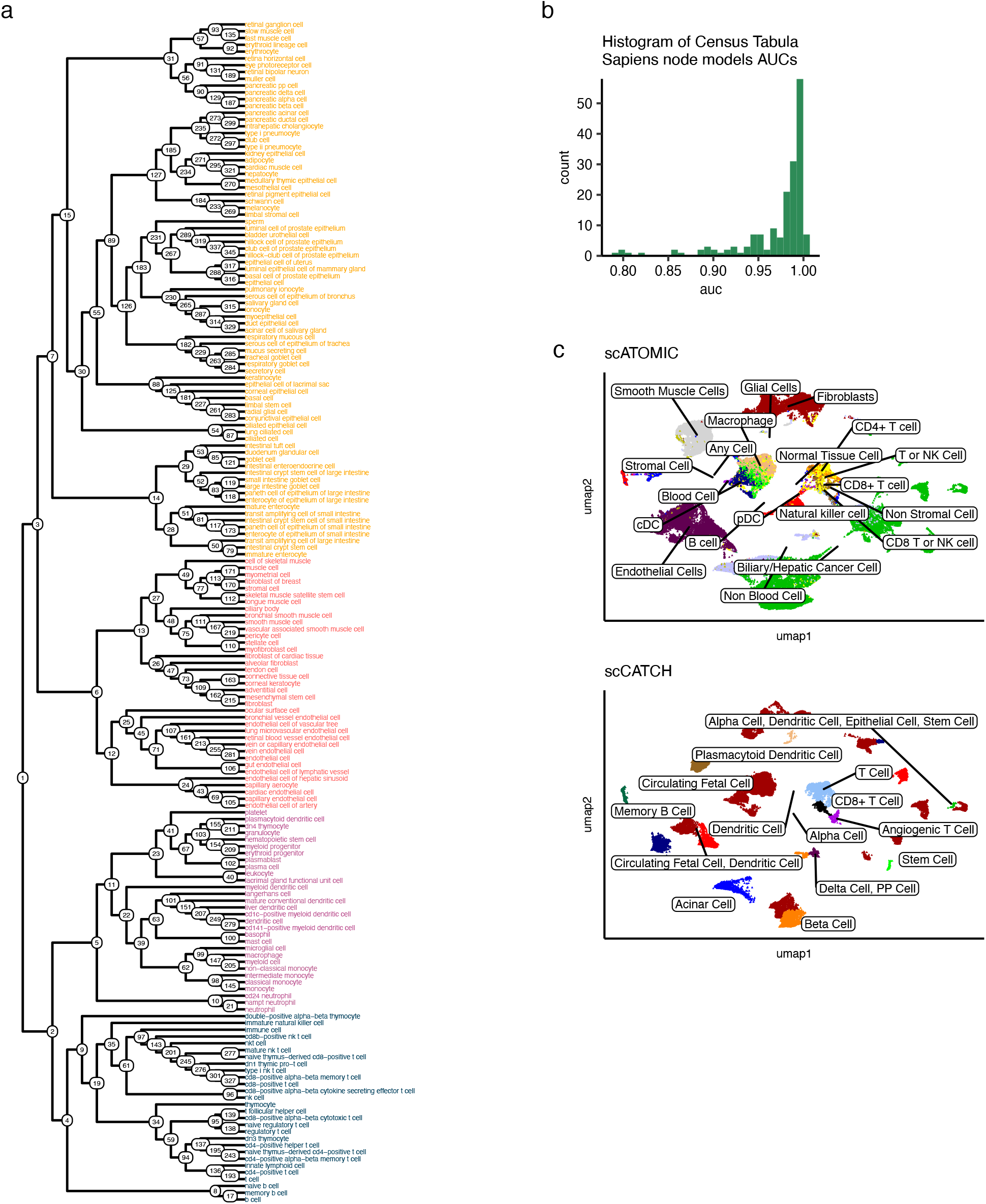
Tabula Sapiens cell-type hierarchy and benchmarking data. **a,** Dendrogram plotting the cell-type hierarchy derived from the Tabula Sapiens. Cells are colored by major compartment; blue, lymphoid; purple, myeloid; red, stromal; orange, epithelial. **b,** Histogram of receiver operator areas under the curves (AUC) for all nodes during Tabula Sapiens model training. **c,** Uniform manifold approximation and projection (UMAP) plots of 57,530 cells from a pancreatic cancer study colored by predicted cell-type with overlaid labels.

**Figure S2.**
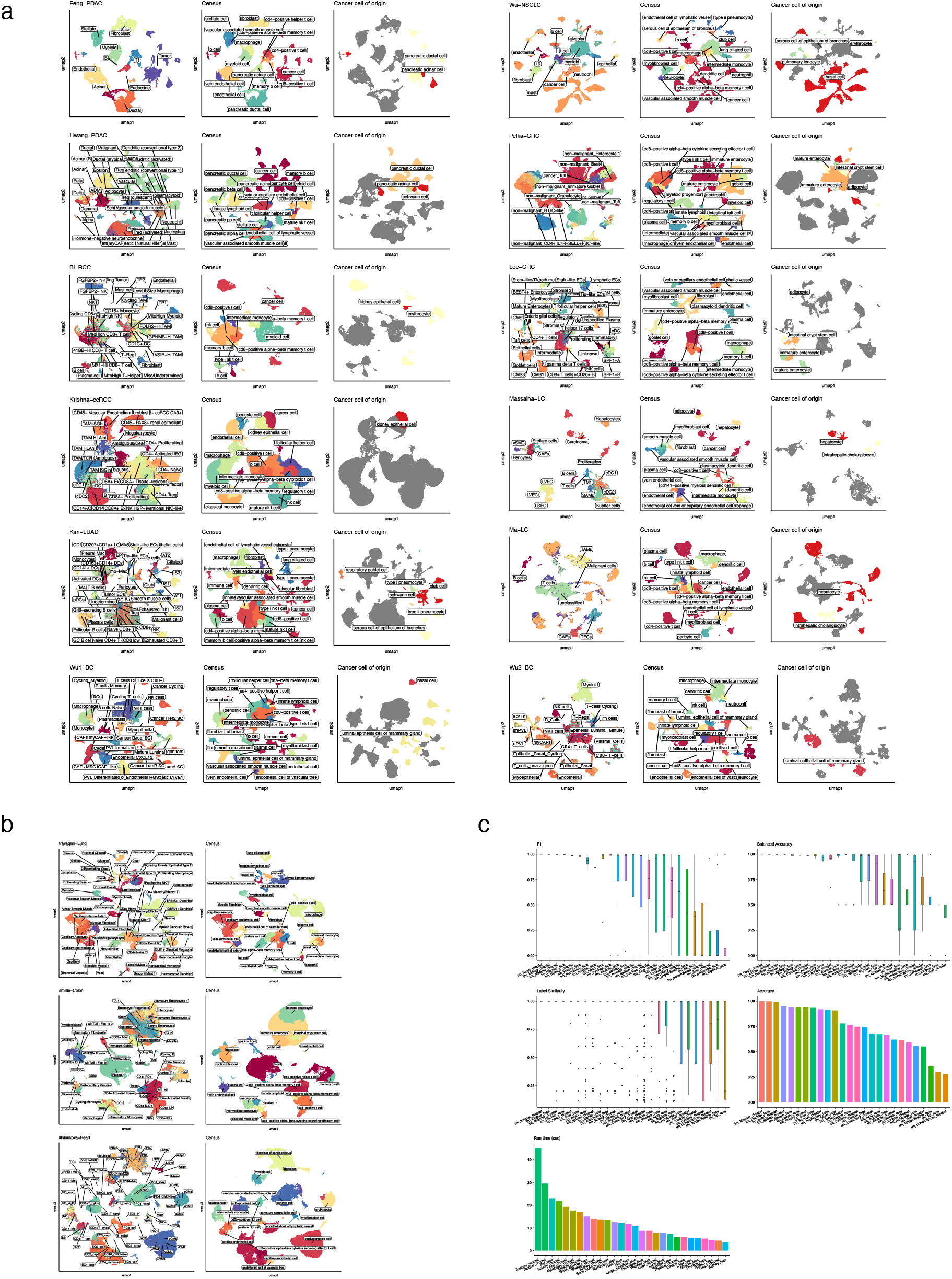
Evaluating Census on human and mouse datasets. **a,** Uniform manifold approximation and projection (UMAP) plots of 12 cancer datasets colored by cell-type annotation with overlaid labels. Left, original author annotations; middle, Census annotation; right, Census predicted cancer cell of origin. PDAC, pancreatic ductal adenocarcinoma; RCC, renal cell carcinoma; ccRCC, clear cell renal cell carcinoma; LUAD, lung adenocarcinoma; BC, breast cancer; NSCLC, non-small cell lung cancer; CRC, colorectal cancer; LC, liver cancer. **b,** UMAP plots of Census annotations on normal human lung, colon, and heart tissues colored by cell-type and with overlaid labels. Left, original author annotations; right, Census annotation. **c,** Evaluating Census prediction performance on tissues from the Tabula Muris. Boxplots show median (line), 25^th^ and 75^th^ percentiles (b ox) and 1.5xIQR (whiskers). Points represent outliers.

